# Stoichiometric homeostasis of wetland vegetation along a nutrient gradient in a subtropical wetland. Understanding stoichiometric mechanisms of nutrient retention in wetland macrophytes

**DOI:** 10.1101/221465

**Authors:** Paul Julian, Stefan Gerber, Rupesh K. Bhomia, Jill King, Todd Z. Osborne, Alan L. Wright

## Abstract

Nutrient homeostasis relates ambient stoichiometric conditions in an environment to the stoichiometry of living entities of the ecosystem. In wetland ecosystems, vegetation can be a large, highly variable and dynamic sink of nutrients. This study investigated stoichiometric homeostasis of dominant emergent and submerged aquatic vegetation (EAV and SAV, respectively) within two treatment flow-ways (FW) of Everglades Stormwater Treatment Area 2 (STA-2). These FW encompass a large gradient in plant nutrient availability. The hypotheses of this study is that wetland vegetation is non-homeostatic relative to ambient nutrients and consequently nutrient resorption will not vary along the nutrient gradient. We developed a framework to investigate how vegetation uptake and resorption of nutrients contribute separately to homeostasis. Overall, the wetland vegetation in this study was non-homeostatic with respect to differential uptake of nitrogen (N) vs. phosphorus (P). Resorption evaluated for EAV was high for P and moderate for N, resorption efficiency did not significantly vary along the gradient and therefore did not affect overall homeostatic status. Nutrient addition experiments may help to compensate for some of the limitation of our study, especially with respect to resolving the primary nutrient source (organic vs. inorganic sources, water vs. soil compartment) and nutrient utilization rates.

---

> “*The environment not only determines the conditions under*
>
> *which life exists, but the organisms influence the*
>
> *conditions prevailing in their environment*.”
>
> — - (Redfield 1958)

A tenet of ecological stoichiometry is that abundance of carbon (C), nitrogen (N), phosphorus (P) or other elements is regulated by reciprocal interaction between organisms and their environment (Redfield 1958), and this relationship forms the underlying framework of stoichiometric homeostasis. In homeostatic organisms, nutrient stoichiometry is relatively fixed while non-homeostatic organisms display flexibility with their respective nutrient stoichiometry. Homeostatic organisms regulate their elemental composition by adjusting bulk material uptake (i.e. ingestion rate), degree of food selectivity for growth and post-assimilation processes to regulate the acquisition of elements. Elemental uptake by non-homeostatic organisms, is mostly controlled by supply (i.e. nutrient availability) rather than demand (i.e. relative growth rate) with limits to nutrient loss processes such as excretion in animals and senescence in plants (Killingbeck 1996; McGroddy et al. 2004).

The concept of stoichiometric homeostasis was formulated as a means to understand patterns of C, N and P in zooplankton to evaluate food quality and nutrient recycling (Jotaro Urabe et al. 1995; Elser and Urabe 1999; J. Urabe and Sterner 2001; Sterner and Elser 2002; Malzahn et al. 2007; Mulder 2007; Mulder and Bowden 2007). Later, the concept has been broadened to understanding food web stoichiometric dynamic (Persson et al. 2010), specific stream food web stoichiometric interactions (Feijoó et al. 2014), resource dependence of bacteria (Makino et al. 2003), and grassland community level responses to climate change (Gu et al. 2017). The stoichiometric homeostasis conceptual model is thus of particular interest with respect to nutrient enrichment and biotic uptake processes due to selective uptake of nutrients during nutrient enriched conditions (Sterner and Elser 2002; Small et al. 2009).

Stoichiometric flexibility or homeostatic plasticity (i.e. non-homeostasis) can manifest across different spatial scales from organisms to ecosystems. At the organismal level, stoichiometric flexibility can occur through changes in nutrient allocation to tissue types or biochemical changes in cellular composition (Sistla and Schimel 2012; Rivas-Ubach et al. 2012). At the ecosystem level, changes in stoichiometry can occur through changes in ecosystem composition including changes in consumer and producer species composition and encroachment of invasive species (Sistla et al. 2015). Therefore, stoichiometric flexibility is driven by species-specific nutrient biomass storage, physiological plasticity, interspecific competition and the variability in nutrient supply versus demand (Elser et al. 2010; Sardans et al. 2012; Sistla et al. 2015).

Heterotrophs (i.e. herbivores) are constantly faced with stoichiometric mismatch but can combat it to maintain constant elemental composition through strategies of selective consumption or elimination of particular elements (Filipiak and Weiner 2014). Meanwhile, autotrophs such as plants have internal mechanisms to continue building up biomass and stoichiometric composition through selective uptake and internal nutrient retention processes such as nutrient resorption. Nutrient resorption, is a process in which nutrients are withdrawn from senescing leaves before abscission to be used for new growth or stored for later use (Van Heerwaarden et al. 2003). It is assumed to be one of the most important strategy used by plants to conserve nutrients (Rejmánková 2005).

Most nutrient resorption studies focus on terrestrial (upland) evergreen, deciduous, forb and graminoid species (Killingbeck 1996; Aerts and Chapin 1999; Van Heerwaarden et al. 2003; Peng et al. 2016) with only few studies investigating wetland plant species (Miao 2004; Rejmánková 2005; Rejmánková and Snyder 2008). Rejmánková (2005) observed relatively high nutrient resorption efficiencies for wetland plant species including cattail (*Typha domingensis*) for both N and P across a variety of nutrient limiting conditions. In a nutrient addition study, Rejmánková and Snyder (2008) observed high N and P resorption across P-limiting conditions in cattail and spike rush (*Eleocharis* spp.). Meanwhile, Miao (2004) observed high P and low N resorption in two wetland macrophytes, sawgrass (*Cladium jamaicense*) and cattail in P-limiting conditions. These studies suggest that P resorption is typically high regardless of P limitation status while N resorption can vary depending on N availability and can contribute significantly to stoichiometric homeostatic.

While limited to a few studies, prior work suggest that plants range in their degree of homeostasis from very weak to non-homeostatic (Yu et al. 2011; Gu et al. 2017). However, these studies focused on terrestrial grasses and shrub species and not wetland plant species distributed along a nutrient gradient. In contrast to upland ecosystems biogeochemical processes in wetlands are influenced by specific non-biotic (inundation, geochemical precipitation of nutrients) and biotic (i.e. plant, periphyton, algae, etc.) interactions that can alter the oxidation-reduction and pH environment (Reddy and DeLaune 2008; Kadlec and Wallace 2009). These interactions pose additional challenges to producers with respect to nutrient supply and demand. Earlier work in wetlands shows a decrease of plant P concentration with decreasing ambient P concentration (Richardson et al.,1990, Julian et al., 2019). This suggests non-homeostatic behavior although the author did not test for homeostasis. Therefore, the first objective of this study was to evaluate the strength of stoichiometric homeostasis in dominant wetland species along a nutrient gradient in two flow-ways (FW) within the Everglades Stormwater Treatment Areas (STAs; Fig. 1). These treatment wetlands use emergent and submerged aquatic vegetation (EAV and SAV, respectively) to remove nutrients via biotic uptake. Prior studies of nutrient resorption with respect to emergent aquatic vegetation (EAV) suggest that EAV species use both “conservation of use” and “enhanced acquisition” of P to fulfill nutrient demand in P-limited systems (Miao 2004; Rejmánková 2005; Rejmánková and Snyder 2008). Moreover these studies indicate that EAV utilize the “conservation of use” strategy making resorption dependent upon nutrient availability. Sterner and Elser (2002) suggest that autotrophic organism’s stoichiometry is not solely dictated by nutrient availability but also biophysical, physiological adaptions and optimal nutrient utilization. The first objective of this study was to determine the degree of stoichiometric homeostasis of wetlands plants along a strong nutrient gradient with the hypothesis that EAV and SAV are generally homeostatic relative to the ambient environment due to biological adaptions in nutrient uptake, utilization and storage. The second objective of this study was to evaluate nutrient resorption dynamics of EAV species within the treatment wetland and with the hypotheses that resorption efficiency (RE) significantly differs as a function of nutrient limitation status (i.e. N or P limitation) and the degree of stoichiometric homeostasis observed among wetland plants is caused by differences in nutrient resorption.

**Figure 1.**
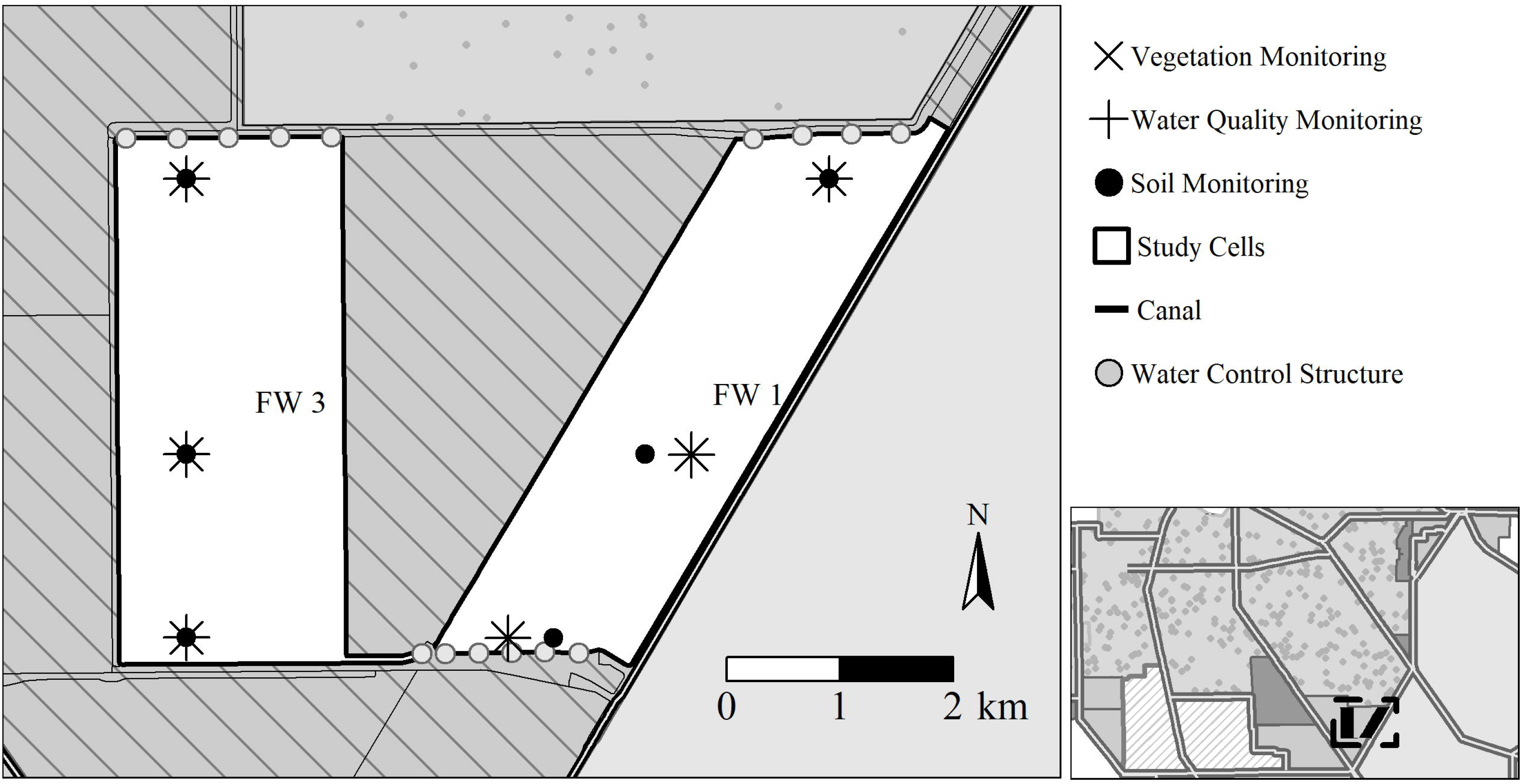
Surface water, soil and vegetation monitoring locations within Everglades Stormwater Treatment Area-2 cells 1 (right) and 3 (left). Cell 1 is predominately emergent vegetation and Cell 3 is predominately submerged aquatic vegetation. Operationally these cells are identified as flow-way 1 and 3, respectively.

## Methods

### Study Area

A total of six Everglades STAs with an approximate area of 231 km^2^ are located south of Lake Okeechobee in the southern portion of the Everglades Agricultural Area (Fig. 1). The Everglades STAs are established on land which earlier had natural wetlands and/or agricultural farmlands under sugarcane production. The primary source of inflow water to the STAs is agricultural runoff originating from approximately 284 km^2^ of farmland upstream. Everglades STA treatment cells or FWs are comprised of a mixture of EAV and SAV communities in several configurations including EAV and SAV treatment cells arranged in parallel or in series (Chen et al. 2015).

Stormwater Treatment Area-2 (STA-2) has been in operation since June 1999 with an effective treatment area of approximately 63 km^2^ divided into eight treatment cells. This study was conducted in two cells, FWs 1 and 3. The vegetative community of FW 1 is comprised predominately of EAV vegetation including *Typha domingensis* Pers. (cattail) and *Cladium jamaicense* Crantz (sawgrass) while FW 3 is dominantly SAV including *Chara* spp. (muskgrass), *Potamogeton* spp. (pondweed) and *Najas guadalupensis* Spreng (southern naiad) with approximately a quarter of the FW occupied by EAV species. Furthermore, prior to STA-2 construction FW 1 was a historic natural wetland while approximately three-quarters of FW 3 was previously farmed and is now managed as an SAV system (Juston and DeBusk 2006).

### Data Source

Data used in this study were collected by South Florida Water Management District (SFWMD) and University of Florida as a part of a larger project within the overall SFWMD’s Restoration Strategies Science Plan to evaluate performance and optimization of the Everglades STAs (South Florida Water Management District 2013). Data from the Science Plan study: Evaluation of P Sources, Forms, Flux and Transformation Processes in the STAs was used for this investigation (UF-WBL 2017; Villapando and King 2018). We used weekly surface water grab samples collected at monitoring locations along a nutrient gradient from inflow to outflow within FWs 1 and 3 in STA-2 to characterize changes in nutrient concentrations during prescribed/semi-managed flow events (Fig. 1). Samples were collected just under the water surface at 10-15 cm depth during prescribed flow events that cycled through various flow/no-flow sequences for FWs 1 and 3. For this study we used the water column parameters total P (TP), total N (TN), and dissolved organic C (DOC) (Villapando and King 2018). In the Everglades system, the surface water organic C pool is predominately composed of the DOC fraction with low particulate OC concentrations (Julian et al. 2017), therefore total OC is assumed to be equivalent to DOC concentrations. We also made use of soil samples collected along the flow transects twice during the dry and wet seasons between 2015 and 2016 and analyzed for TP, TN, and TC. Vegetation tissue were collected from live and senescent above-ground biomass (AGB) vegetation in FW 1 while only live-AGB was sampled from FW 3 at the end of the 2015 (November 2015) and 2016 (September 2016) wet seasons. The ability to collect senescent SAV is extremely difficult as it decomposes rapidly and becomes part of the “litter” immediately, therefore no samples were available for analysis. Vegetation samples were collected from inflow, mid and outflow regions of the FWs within close proximity to the surface water and soil monitoring locations (Fig. 1). Only co-located surface water, soil and vegetation sampling locations were considered for this study (Fig. 1). Nutrient content of vegetation was determined from dried ground and homogenized samples. For conducting analyses and summary statistics, data reported as less than method detection limit (MDL; Table S1) were assigned a value of one-half the MDL, unless otherwise noted.

### Data Analysis

We categorized foliar TN:TP molar ratios for EAV along the FW 1 transect into potential N-limitation (molar TN:TP<14), potential N and P co-limitation (14 < TN:TP < 16) and P limitation (TN:TP > 16). For SAV along the FW 3 transect we categorized foliar TN:TP molar ratios into potential N-limitation (TN:TP < 27) and potential P-limitation (TN:TP > 27). These potential nutrient limiting categories were consistent with other EAV studies (Koerselman and Meuleman 1996; Rejmánková 2005) and SAV studies (Duarte 1992; Fernández-Aláez et al. 1999). We compared nutrient limitation categories between FW regions (i.e. inflow, mid and outflow) using chi-squared (RxC) analysis.

We evaluated P and N resorption efficiency (RE_TP_ and RE_TN_, respectively) along the FW 1 transect by comparing living (X_GR_) to senescent (X_SEN_) EAV AGB nutrient tissue concentrations. Resorption efficiency was estimated using Eq. 1, consistent with prior studies (Killingbeck 1996; Reed et al. 2012).

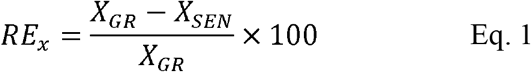

We compared RE_TP_ and RE_TN_ values between P-limiting categories identified above for aquatic vegetation type specific foliar TN:TP molar ratios using the Kruskal-Wallis rank sum test. Resorption efficiency values for each site was compared with fractional distance from inflow using Spearman’s rank correlation. We also compared RE_TP_ and RE_TN_ between stations using Dunn’s multiple comparison rank sum test (‘dunn.test’ R package; Dinno 2015).

We characterized the strength of vegetation TN:TP total stoichiometric homeostasis (H_T_) using the homeostasis coefficient (Eq. 2). Equation 2a is a log-linear model where *y* is the live-AGB TN:TP molar ratio, *x* is the TN:TP molar ratio of the resource, *c* is a constant that determines the intercept and 1/*H*_*T*_ is the slope of the model between the log transformed ratios *x* and *y* and represents the strength of homeostatic maintenance of nutrient ratios within the consumer biomass. Reorganizing Eq. 2a to solve fore 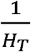 get Eq. 2b.

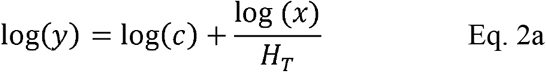

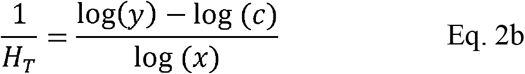

Hypothetically, values of 1/*H*_*T*_ range between zero and one, with values approaching one indicating a non-homeostatic relationship and values approaching zero signifying consumer-resource homeostasis. The degree of homeostasis was defined consistent with prior studies (Makino et al. 2003; Persson et al. 2010; Feijoó et al. 2014) as the following:

1. 0.00 < 1/*H*_*T*_ < 0.25 : Homeostatic
2. 0.25 < 1/*H*_*T*_ < 0.50 : Weakly-Homeostatic
3. 0.50 < 1/*H*_*T*_ < 0.75 : Weakly Non-Homeostatic
4. 0.75 < 1/*H*_*T*_ < 1.00 : Non-Homeostatic

The stoichiometric homeostasis coefficient is an integrated metric to relate the ambient environment’s stoichiometry to that of the vegetation biomass. Moreover H_T_ can be used to determine if the observed variation in the biota stoichiometry is the result of physiological adjustments or reflects species-specific turnover (Elser et al. 2010). To assess the homeostatic relationship of the vegetation relative to the ambient environment, N:P ratios of resource pools were compared to the N:P ratios of AGB for each vegetation community. The communities’ uptake pathways differ between SAV and EAV - SAV primarily assimilate nutrients through the water column while EAV obtain nutrients from the soils via soil solution (Richardson and Marshall 1986; Reddy et al. 1999; Dierberg et al. 2002). Therefore, we compared AGB N:P values of both EAV and SAV to surface water and soil N:P values (denoted as x in Eq. 2). Due to offset sampling times, soil and AGB data were paired only when samples were collected within six months of each other for each site. Site-specific surface water mean TN:TP values during this six-month timeframe were paired with AGB values. Biota TN:TP molar ratios were identified as each respective vegetation type live-AGB TN:TP molar ratio (y in Eq. 2). We used the Theil-Sen single median linear model (‘mblm’ package; Komsta 2013) to evaluate biota and resource TN:TP molar ratios to estimate c and 1/H_T_ in Eq. 2.

To evaluate the overall homeostatic versus non-homeostatic status among vegetation within a FW, we evaluated the slope of the mblm linear models for each FW to determine if the slope was significantly different from 0.50 using the One-Sample Wilcoxon rank signed test. This test is similar to the t-test slope comparison method identified by Zar (2010) for parametric linear models. The 0.50 threshold was selected as it is the criterion that differentiates homeostasis from non-homeostasis (see above).

We also characterized the degree of stoichiometric homeostasis with respect to differential resorption similar to the approach for stoichiometric homeostasis (H_T_) above. Resorption efficiency homeostasis (H_RE_) was evaluated for EAV to assess whether RE changes with respect to the ambient environment. We used mblm to evaluate the relative loss of nutrients after resorption (*y*_*RE*_) relative to resource TN:TP (x) where the slope of linear model is 1/H_RE_. The relative loss of nutrients after resorption (*y*_*RE*_) or the senescence efficiency is represented by Eq. 3. In the supplementary material we develop the framework to evaluate how resorption contributes to homeostasis, the resulting relationship is given in Eq. 4 which is very similar to Eq. 2. The subtraction of one on the left-hand side in Eq. 4 originates from the fact that this equation states the vegetation’s effort into homeostasis via resorption, where zero would yield an effort to maintain full homeostasis and negative one would mean that there is no homeostasis with respect to resorption (Supplemental Material). The regression constant c is different than in Eq. 2, but is in both cases irrelevant to the calculation of stoichiometric homeostasis.

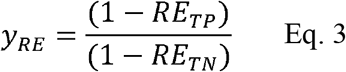

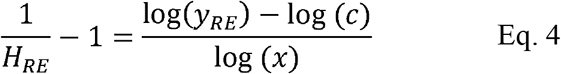

In the supplemental material, we have developed a framework which can relate the two major pathways for homeostasis (resorption and uptake) with each other. Total homeostasis (*H*_*T*_; Eq. 2) and the vegetation’s effort to maintain homeostasis can then be partitioned into contributions from resorption (Eq. 4) and uptake:

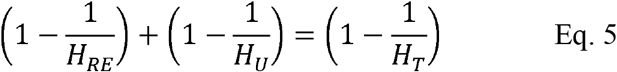

All statistical operations were performed with R© (Ver 3.1.2, R Foundation for Statistical Computing, Vienna Austria), unless otherwise stated all statistical operations were performed using the base R library. The critical level of significance was set at α=0.05.

## Results

Surface water DOC, TN concentrations, DOC:TP and TN:TP molar ratios were overall higher in FW 3 (SAV) while surface water TP concentrations and DOC:TN molar ratios were higher in FW 1 (EAV) (Table 1). Soil organic matter, as indicated by loss-on-ignition, was greater in FW1 while soil Ca concentrations were greatest in FW3 soils indicating that FW1 soil is more organic (i.e. peat) and FW3 more mineral (i.e. marl). In addition to qualitative differences between soil composition, FW1 soils were generally more enriched with TP and TN. Among carbon and nutrients, soil TP had the highest coefficient of variation between and within sites and FWs with an overall coefficient of variation of 48%. Overall, FW 1 exhibited the highest coefficient of variation for soil TP with 40% spatial and temporal variability (Table 1).

**Table 1.**
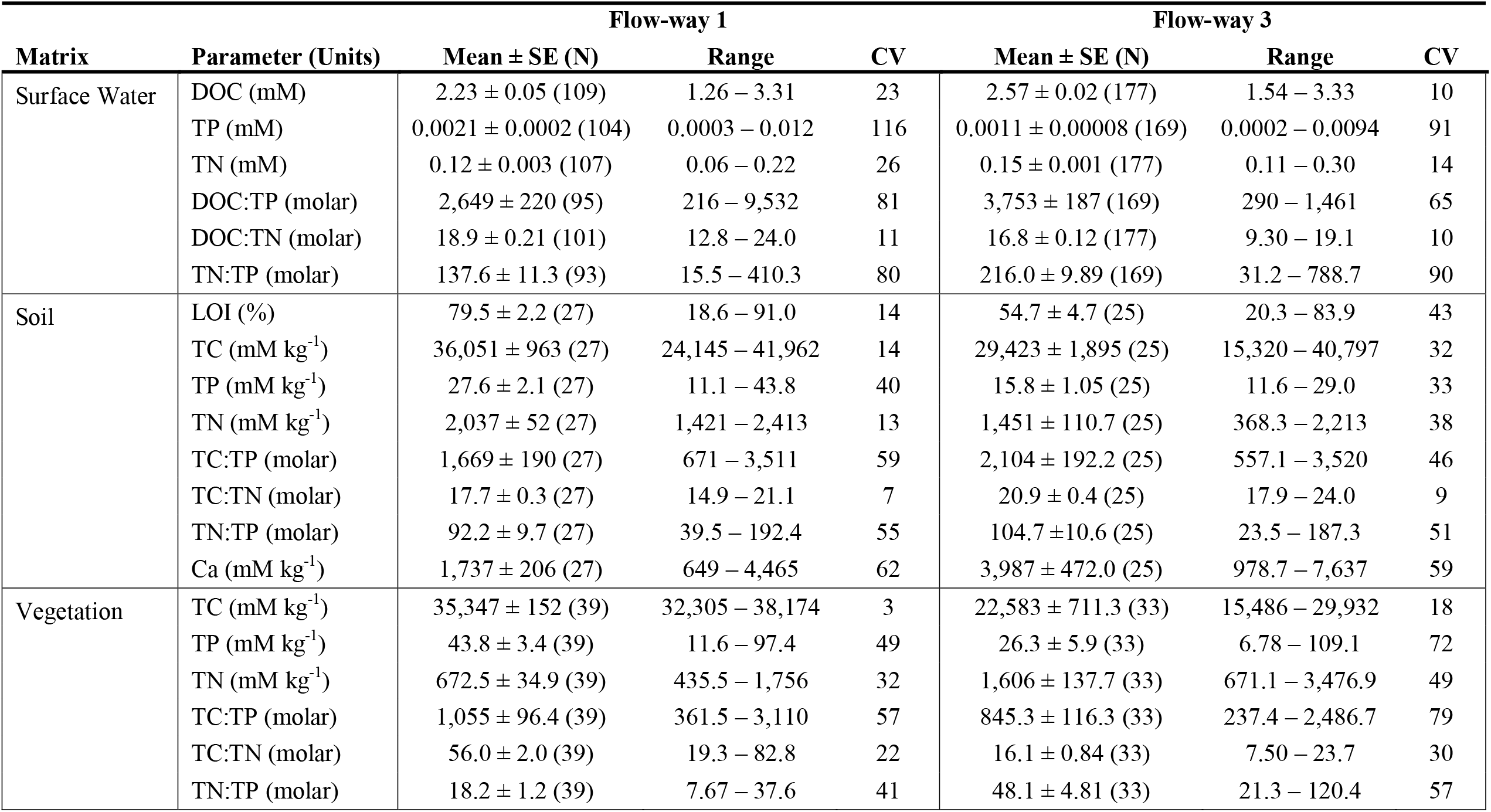
Summary statistics for variables and matrices used in this study of samples collected along the flow-way (FW) 1 and 3 transect within Stormwater Treatment Area-2. Summary statistics include mean, standard error (SE), sample size (N), range, and coefficient of variation (CV). Matrices include surface water, recently accreted soil and live aboveground biomass of sampled vegetation Note: Dissolved Organic Carbon = DOC; Total Phosphorus = TP; Total Nitrogen = TN; Total Carbon - TC; Loss-on-ignition = LOI; Calcium = Ca.

### Tissue Stoichiometry

Live AGB TN : TP molar ratios were greatest in FW 3 (Table 1 and Fig. 2). Vegetation from both FWs spanned their respective potential N and P-limiting conditions with most values above the P limits identified for EAV (TN:TP>16) and SAV (TN:TP>27) communities (Table 1 and Fig. 2). Furthermore, AGB TC:TN values were greater in FW 1, ranging from 19.3 to 82.8 whereas FW 3 ranged from 7.5 to 23.7 suggesting the potential for N-limitation in FW 1 (Fig. 2). AGB TN:TP values were significantly different along the FW 1 transect (χ^2^=19.4, df=4, ρ<0.01) characterized by strong nutrient decline along the FW from inflow to outflow. Several values below the TN:TP value of 14 (indicating N limiting conditions) were observed, most of which occurred at the Inflow and Mid regions of the FWs and all values above the P-limiting criteria occurred in the Outflow region. Meanwhile, nutrient limiting conditions were not significantly different along the FW 3 flow path (χ^2^=5.6, df=2, ρ=0.06) with the majority of the TN:TP values greater than 27 indicating potential P-limitation, and four samples less than 27 occurring in the Inflow and Mid regions of FW.

**Figure 2.**
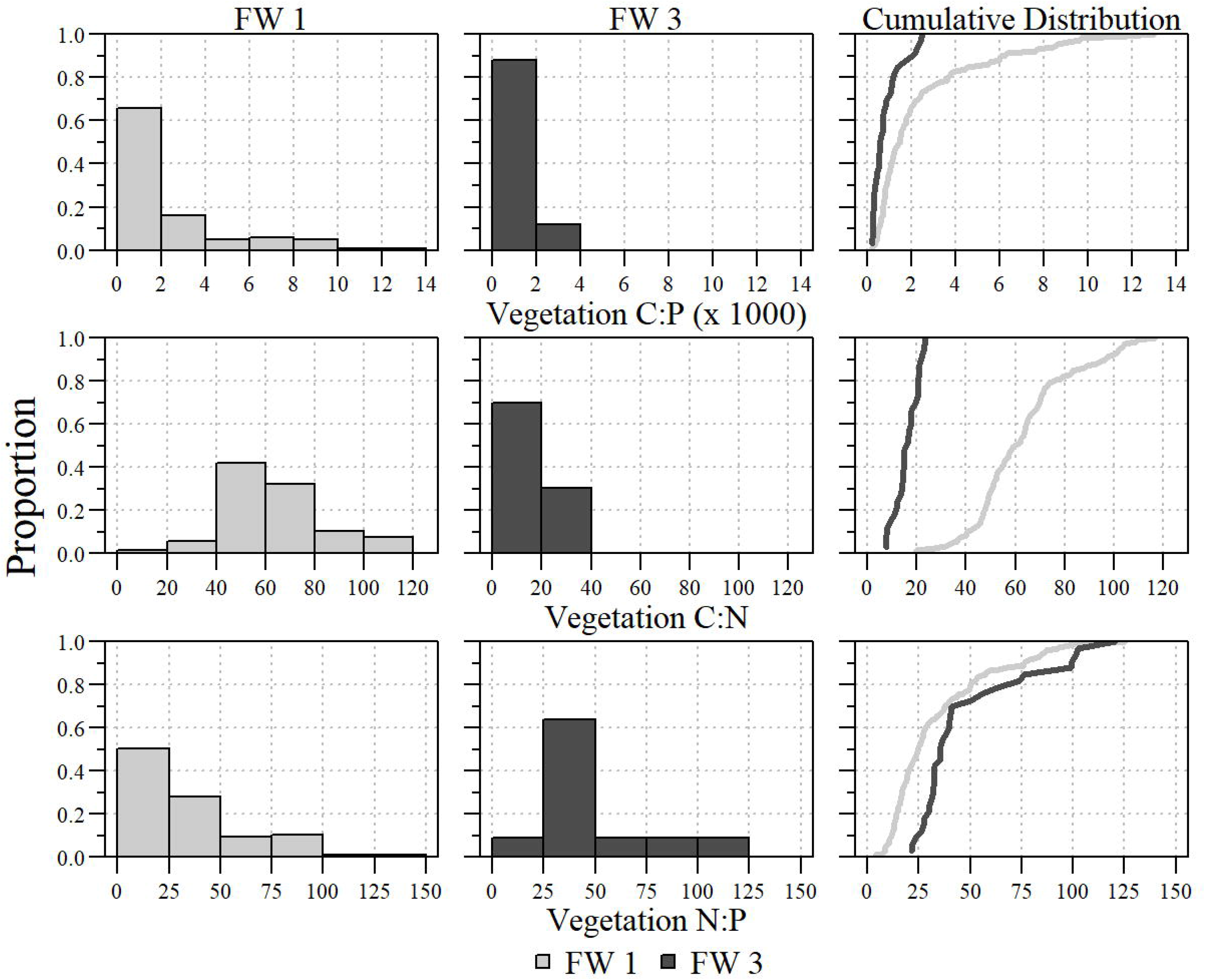
Distribution of plant foliar stoichiometric relationships within flow-way (FW) 1 and FW 3 of Stormwater Treatment Area-2. Left and middle panels are histogram representations of each respective stoichiometric relationship. The right panel is a cumulative distribution of stoichiometric relationship for each FW.

### Stoichiometric Homeostasis

Changes in AGB stoichiometry occurred concurrent with changes in both surface water and soil resource stoichiometry pools (Fig. 3). Log-log models comparing vegetation N:P to resource N:P resulted in slopes significantly different from zero (Table 2) with varying levels of fit. The model with the lowest R^2^ was the SAV (FW 3) N:P to surface water N:P (Table 2). The relationship between the vegetation and the ambient environment depends on which resource pool was considered and thereby where each vegetation type falls on the H_T_ continuum. For instance, EAV is weakly homeostatic if the water column is considered to be the major nutrient source and non-homeostatic if nutrients are assumed to be taken up from the soil (Fig. 4 and Table 2). The homeostasis coefficients, 1/H_T_, when considering soil as a potential resource pool, were significantly greater than 0.5 for EAV (V=153870, ρ<0.01) and SAV (V=14455, ρ<0.01) indicating that vegetation is non-homeostatic with respect to soil TN:TP (Fig. 4). In fact, the homeostasis coefficient 1/H_T_ for EAV sourcing nutrient from soil is greater than 1. Meanwhile, 1/H_T_ values when considering surface water as a potential resource pool were significantly less than 0.5 for EAV (V=53539, ρ<0.01) and not significantly different for SAV (V=15452, ρ=0.36) suggesting EAV and SAV with respect to surface water is weakly homeostatic to weakly non-homeostatic, respectively (Fig. 4).

**Table 2.** Overall log-log median based model statistics of vegetation nitrogen to phosphorus (N:P) versus soil and surface water resource pools between emergent and submerged aquatic vegetation (EAV and SAV, respectively) communities within Stormwater Treatment Area 2 Flow-way 1 and Flow-way 3, respectively.

| Resource | Community | Slope<br>(1/H<sub>T</sub>) | Intercept<br>(c-value) | R<sup>2</sup> | Median<br>Absolute<br>Deviation | V-statistic | df | p-value |
| --- | --- | --- | --- | --- | --- | --- | --- | --- |
| Soil | EAV | 1.62 | -3.70 | 0.61 | 1.50 | 168032 | 37 | <0.01 |
|  | SAV | 0.66 | 0.84 | 0.72 | 0.47 | 19947 | 22 | <0.01 |
| Water Column | EAV | 0.46 | 0.70 | 0.60 | 0.34 | 116683 | 37 | <0.01 |
|  | SAV | 0.51 | 0.93 | 0.17 | 0.94 | 23162 | 26 | <0.01 |

**Figure 3.**
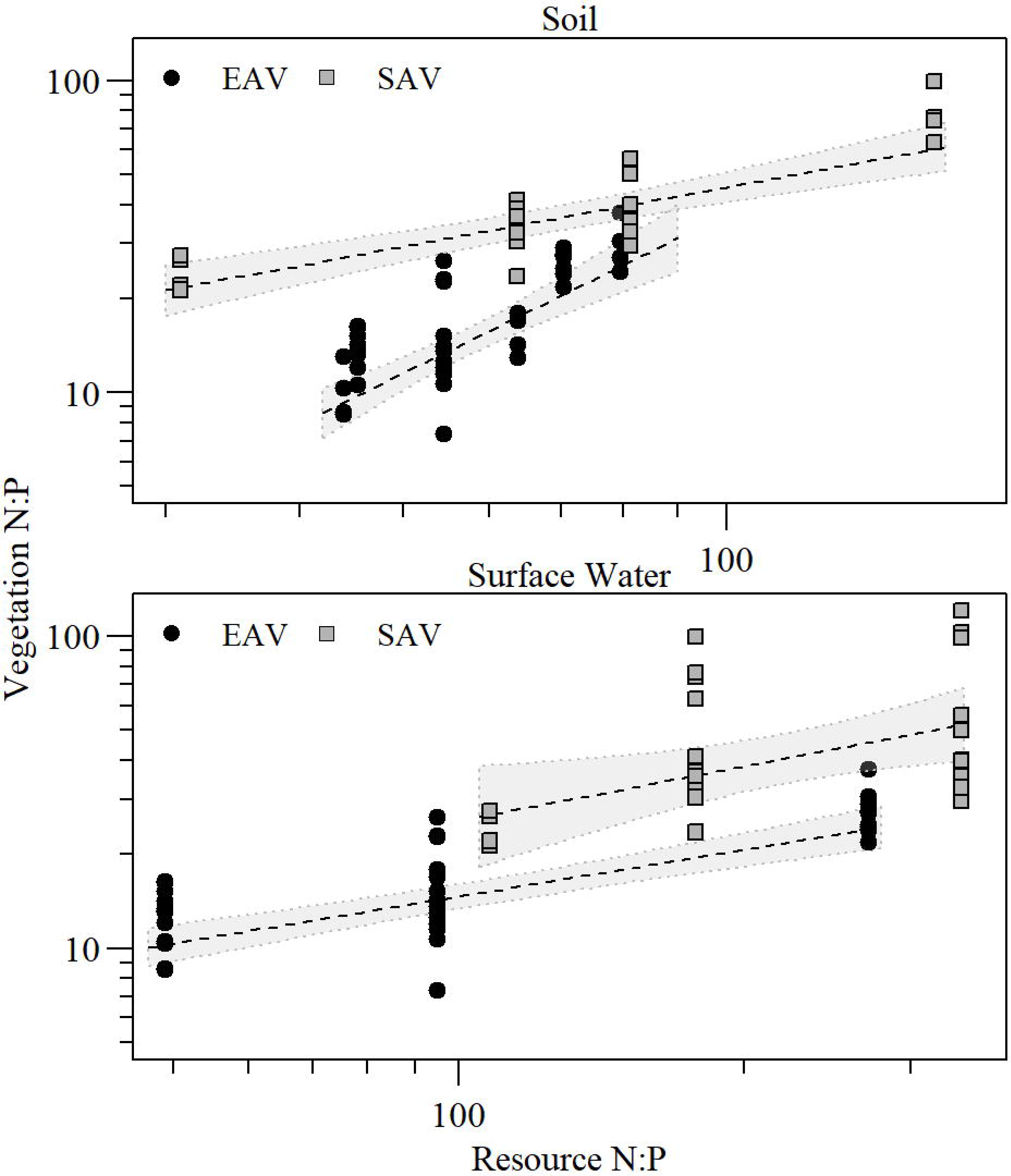
Resource (Top: Soil and Bottom: Surface Water) and vegetation nitrogen to phosphorus molar ratio (N:P) with Theil-Sen single median linear model ± 95% confidence interval for each vegetation type. Note both axes are log-scale.

**Figure 4.**
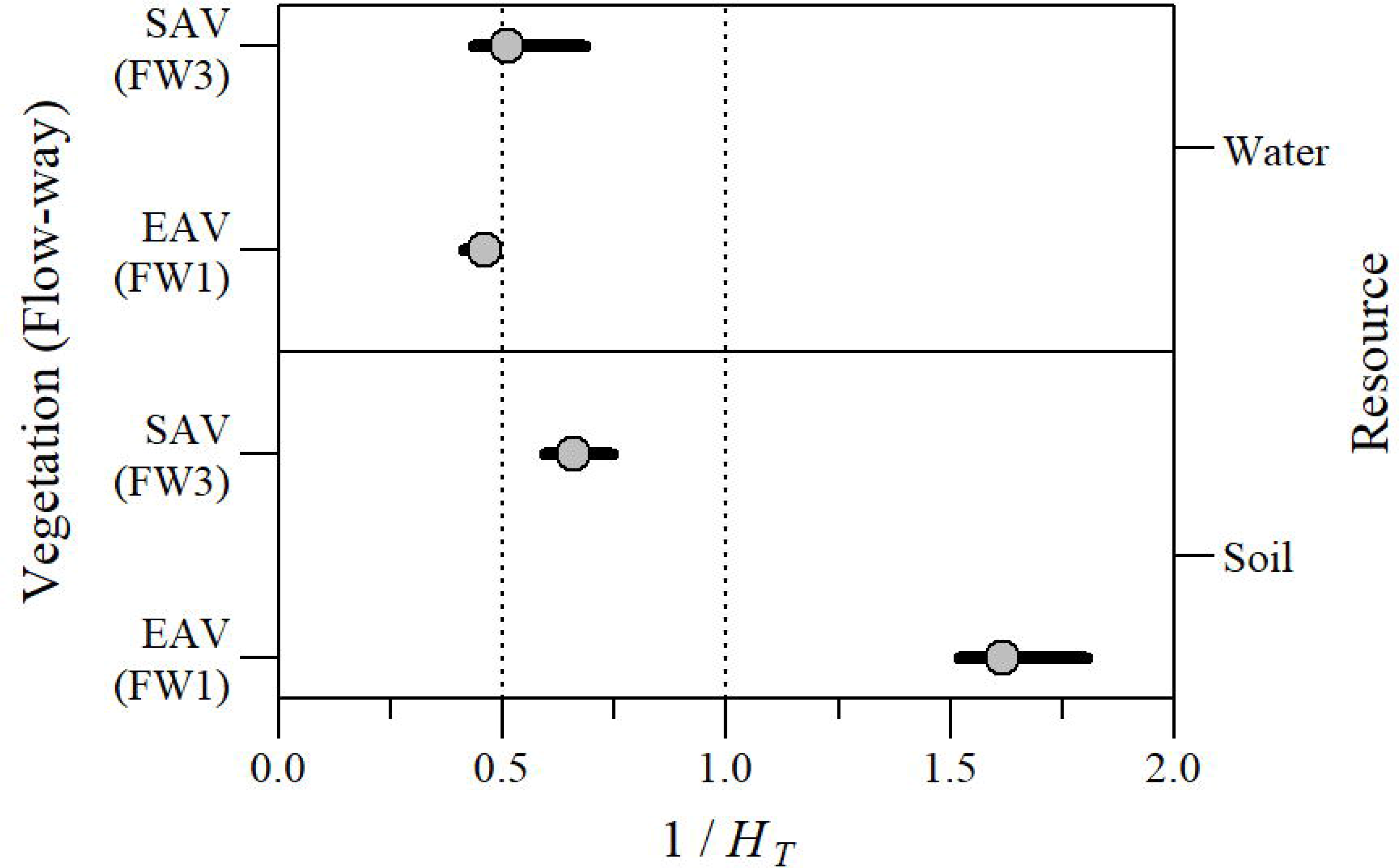
Homeostasis coefficient (1/H_T_) and 95% confidence interval with respect to soil and surface water resources for emergent aquatic vegetation and submerged aquatic vegetation (EAV and SAV, respectively) vegetation communities and flow-way. Dashed line indicates the homeostasis/non-homeostasis threshold (0.5) and the theoretical maximum 1 / H_T_ (i.e. 1.0).

### Resorption Efficiency

A total of 38 plant tissue samples were collected during the vegetation sampling events where both live and senesced AGB were available from EAV (FW1). Cattail comprised most of the species collected (N=36) with one sample each from lily and sawgrass. The estimated mean RE_TP_ value for cattails was 74 ± 2 % (N=36), with the individual RE_TP_ values for water lily and sawgrass being 70 % and 77 %, respectively (Table 3). Along FW 1, cattail RE_TP_ was not significantly correlated with fractional distance downstream along the resource gradient (r_s_=0.21, ρ=0.22) or statistically different between sites (χ^2^= 3.9, df=2, ρ=0.14). The estimated mean RE_TN_ value for cattails was 32 ± 3 % (N=36), with RE_TN_ values for just sawgrass observed to be 2.3% (Table 3). Much like RE_TP_, cattail RE_TN_ was not significantly correlated with fractional distance downstream (r_s_ = −0.04, ρ=0.84) or statistically different between sites (χ^2^= 2.9, df=2, ρ=0.23). Both TP and TN RE did not significantly differ between nutrient limitation categories as indicated by tissue TN:TP molar ratios (χ^2^= 0.74, df=2, ρ=0.70 and χ^2^= 0.38, df=2, ρ=0.83, respectively).

**Table 3.** Resorption efficiency summary statistics for emergent aquatic vegetation species collected within Stormwater Treatment Area 2, FW 1. Total Phosphorus Resorption efficiency (RE_TP_) calculated with equation 1, using both living and senescent living aboveground biomass. Summary statistics include mean, standard error, range, coefficient of variation and sample size (N).

| Species | Total Phosphorus |  |  |  | Total Nitrogen |  |  |  |
| --- | --- | --- | --- | --- | --- | --- | --- | --- |
|  | Mean ± SE (%) | Range (%) | Coefficient of Variation | N | Mean ± SE (%) | Range (%) | Coefficient of Variation | N |
| *Typha domingensis* Pers.<br>(cattail) | 74 ± 2 | 24 – 91 | 20.1 | 36 | 33 ± 2 | 1.8 - 56 | 39.8 | 29 |
| *Nymphaea odorata* Aiton<br>(water lily) | 70 | --- | --- | 1 | --- | --- | --- | --- |
| *Cladium jamaicense* Crantz<br>(sawgrass) | 77 | --- | --- | 1 | 2.3 | --- | --- | 1 |

### Partitioned Contributors to Stoichiometric Homeostasis

Senescence efficiency ranged from 9% to 76% for P and 44% to 98% for N in cattail along FW 1. Therefore, the relative stoichiometric loss of nutrient after resorption (*y*_*RE*_; Eq. 4) ranged from 0.16 to 0.73 for cattail along FW 1. Generally, the relative stoichiometric loss of nutrient after resorption did not vary relative to the resource availability. When considering soil as a potential resource pool relative to stoichiometric resorption, the slope was significantly different from zero (V=33519, ρ<0.05) with a value of 0.24 indicating weakly homeostatic. Additionally, when considering surface water as a potential resource pool the slope of the model was significantly different from zero (V=21674, ρ<0.05) with a value of 0.05, suggesting a homeostatic relationship.

## Discussion

All organisms are faced with the challenges of acquiring sufficient energy and nutrients, with most organisms dealing with a mixture of these resources in the environment that is imbalanced relative to their requirements. Resource constraints and imbalances can alter individual species strategies, population dynamics, inter-specific competition for resources, and even influence key ecosystem processes (Frost et al. 2005). Ultimately, stoichiometric homeostasis is largely controlled by a set of key physiological processes that regulate uptake, incorporation and eventual release of nutrients (Sterner and Elser 2002). These processes must contend with what is available in the ambient environment and can manifest as stoichiometric mismatches and lead to nutritional imbalances. We used the concept of stoichiometric homeostasis to evaluate the degree to which plants in a wetland ecosystem respond to variations in nutrient supply. The dominant features of the variable nutrient supply in our system were both a decreasing P load (i.e. uptake during transport) and a diminishing resource gradient (i.e. soil nutrient concentration) with distance from inflow to outflow of two treatment wetland FWs.

Neither of the two types of vegetation (EAV and SAV) in the two studied FWs evaluated here displayed clear homeostatic behavior, in that N:P stoichiometry of AGB remained independent of the N:P stoichiometry of the resources (i.e. soil or water column nutrient pools). Considering the water column as the resource supply, both EAV and SAV displayed weakly homeostatic to weakly non-homeostatic behavior, respectively. However, if soil is considered as the resource pool, EAV and SAV are considered weakly to strongly non-homeostatic, respectively. Generally, EAV use nutrients from the soil while SAV obtains nutrients from the water column.

Early in the development of stoichiometric homeostasis theory it was hypothesized that photoautotrophs (i.e. algae and plants) were considered to have very weak stoichiometric homeostasis (0.25 < 1/H < 0.50), suggesting that they do not strongly regulate biomass stoichiometry relative to the ambient environment due to luxury uptake of nutrients (Sterner and Elser 2002; Reddy and DeLaune 2008; Elser et al. 2010). Since development of the concept, however, several studies have suggested that plants can range from weakly homeostatic to non-homeostatic (1/H >0.50) (Sterner et al. 1998; Elser et al. 2000; McGroddy et al. 2004; Güsewell 2004). Moreover, Demers et al. (2007) suggested that the relationship between external nutrient concentrations (i.e. soil and water column) and foliar tissue nutrient concentrations in aquatic macrophytes indicates a weakly non-homeostatic to non-homeostatic relationship similar to other terrestrial and algae species. This conclusion was confirmed in our study from treatment wetlands of south Florida which received high inputs of P as agricultural runoff.

A very important aspect of stoichiometric homeostasis is the proper pairing of resource to organism along a trophic chain as this affects the efficiency of trophic transfer of. Theoretically, 1/*H* values range from zero to one. However, values can exceed one, and in this study, a homeostasis value greater than one was observed when comparing soil and EAV N:P. This is manifested by a disproportionate increase in AGB N:P with increasing soil N:P. In a review of stoichiometric homeostasis, Persson et al. (2010) reported several studies with 1/*H* values greater than one. As suggested by Smalley et al. (2003), it is possible that non-inhibitory light levels leading to high C-fixation and greater biosynthetic need for N and P to use the additional C could lead to 1/*H*_*N:P*_ > 1. EAV, and especially cattails, are generally not growth-limited by light due to its tall structure above the water and broad leaves. Therefore, it is possible that EAV can respond by disproportionally directing N use towards building other cellular machinery to optimize nutrient use and functionality. An alternative mechanism may be that, in this P-limited system, P acquisition becomes increasingly difficult as N:P increases because P is stored in increasingly inaccessible forms within the resource pool.

It is possible that both vegetation types (EAV and SAV) take up nutrients from soil and water. For EAV, homeostatic behavior appears stronger if the nutrient resource is assumed to come from the soil rather than water column (Fig. 4). By the same token, homeostasis for SAV is stronger if nutrients are assumed to be supplied from the water column (Fig. 3), although the difference between the homeostasis coefficients for SAVs is decidedly lower compared to EAVs (Fig 6). This may in turn partially be explained by lower variability in N:P ratios in EAV compared to SAV, which in turn suggests less flexible N:P ratios and thus stronger homeostasis in EAV.

As expected, stoichiometric mismatches were observed within our study system where resource (i.e. soil or water column) N:P were greater than that of the AGB N:P (Table 1). Both soil and water resource pools differed between FWs with water having a greater variability than that of soil. This is notable as a strong soil P gradient is apparent along the FW (Osborne et al, *In Prep*), potentially indicating changes in P-availability and the physiological limitation to use the existing P-pool. This limitation has been noted in prior studies where uptake is lower in treatments with higher available P (Shaver and Melillo 1984; Craft et al. 1995). Variability and differences in the relative ratio of resources to consumer indicate that vegetation types access available nutrient pools differently. In lake ecosystems, Rattray et al. (1991) determined that SAV take up nutrients from the water column. However, other studies have demonstrated that, while SAV are involved in active uptake of nutrient from the water column, it also modifies the physical environment through photosynthesis and respiration causing the precipitation of P to mineral bound P compounds (Dierberg et al. 2002; Reddy and DeLaune 2008). Submerged macrophyte systems invoke P mineralization by inadvertently manipulating C (via dissolved inorganic C) and pH through respiration and photosynthesis. In FW 3, pH values ranged from 6.9 to 10.4 (Julian et al. 2019) and, combined with a high mineral content environment, can lead to coprecipitation of P with CaCO_3_ (Boyer and Wheeler 1989)(Table 1). Much like their emergent relatives, rooted SAV can also obtain nutrients from the soil but to a lesser degree (Brix 2003; Dhote and Dixit 2009). Stoichiometric mismatch in producer-consumer (i.e. detritus-invertebrate) systems, typically identify nutritional deficiency in food sources (Filipiak and Weiner 2014, 2017; Filipiak 2016). In this case stoichiometric mismatch is indicative less of nutritional deficiency but more of limitation in nutrient utilization and availability as well as the potential for plant-environment nutrient feedbacks discussed above. Regardless of the stoichiometric mismatch observed, we still observe predominantly non-homeostatic responses by vegetation across N to P-limiting conditions for both FWs despite being predominant P.

Generally, plant species respond to changes in nutrient available nutrients through the storage and recycling of nutrients, ultimately shifting absolute and relative tissue nutrient concentrations in a way suggesting a non-homeostatic relationship with their ambient environment (Güsewell and Koerselman 2002; Güsewell et al. 2003; Rejmánková et al. 2008). The resorption of nutrients from senescing tissues is often considered a strategy in nutrient-limiting conditions to conserve scarce nutrients (Aerts and Chapin 1999; Rejmánková et al. 2008). Nutrient resorption efficiency of EAV observed in this study (Table 3) (Table 1) is consistent with values observed for aquatic macrophytes by studies elsewhere (Miao 2004; Rejmánková 2005). Rejmánková (2005) evaluated resorption of macrophyte species from several wetlands across a broad spatial distribution and observed that cattail achieved a maximum RE_TP_ of approximately 80% while the study average was 70%. Rejmánková (2005) also observed that RE_TP_ was significantly greater in P-limiting conditions. Meanwhile, Miao (2004) evaluated RE_TP_ of two aquatic macrophytes in a mesocosm setting with two-nutrient treatment regimens broadly defined as enriched (high nutrient availability) and unenriched (low nutrient availability). Miao (2004) observed RE_TP_ ranging from 69 to 82% for sawgrass and 61 to 71% for cattail between treatments with no treatment effect, indicating that, despite available nutrient conditions, resorption remains relatively high. Miao (2004) also observed RE_TN_ values ranging from 33 to 45% between cattail and sawgrass species with no significant differences between treatment or species. In our study, RE_TP_ or RE_TN_ was not significantly different along the flow way despite potential shifts in relative N or P-limitation as indicated by foliar TN:TP values (Fig. 2) and nutrient stoichiometry of other compartments (Julian et al. 2019). Mean RE_TP_ values observed at the inflow region, where P resources are especially high, were slightly lower with more variability than middle and outflow regions (Fig. 5), potentially suggesting some adaptation. We note that leaching of senesced litter may overestimate active resorption calculated here and in previous studies.

**Figure 5.**
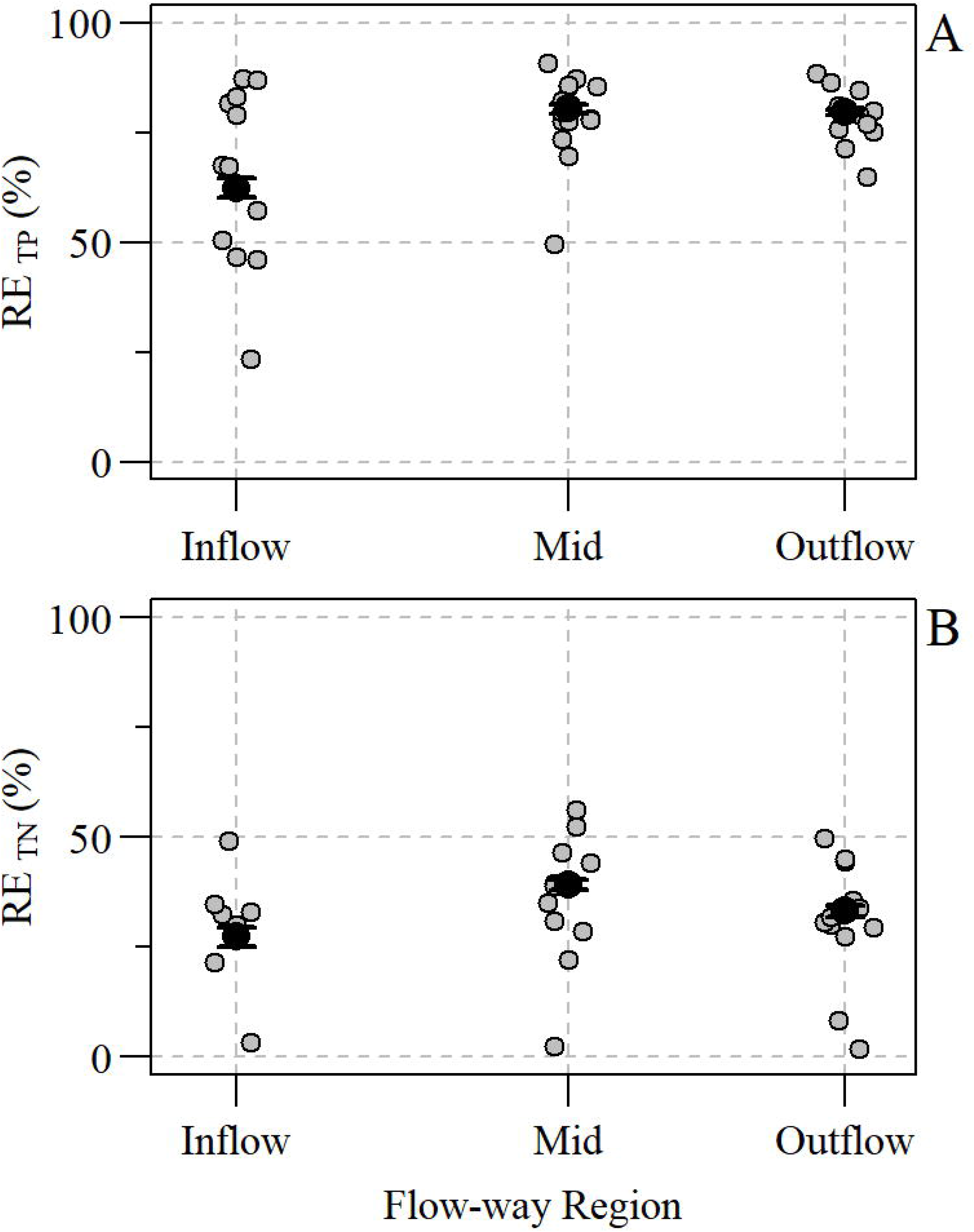
Strip plot of (A) total phosphorus resorption efficiency (RE_TP_) and (B) total nitrogen resorption efficiency (RE_TN_) of cattail within Stormwater Treatment Area-2 flow-way 1. Raw data are represented by grey points, while mean ± standard error are represented by black points.

**Figure 6.**
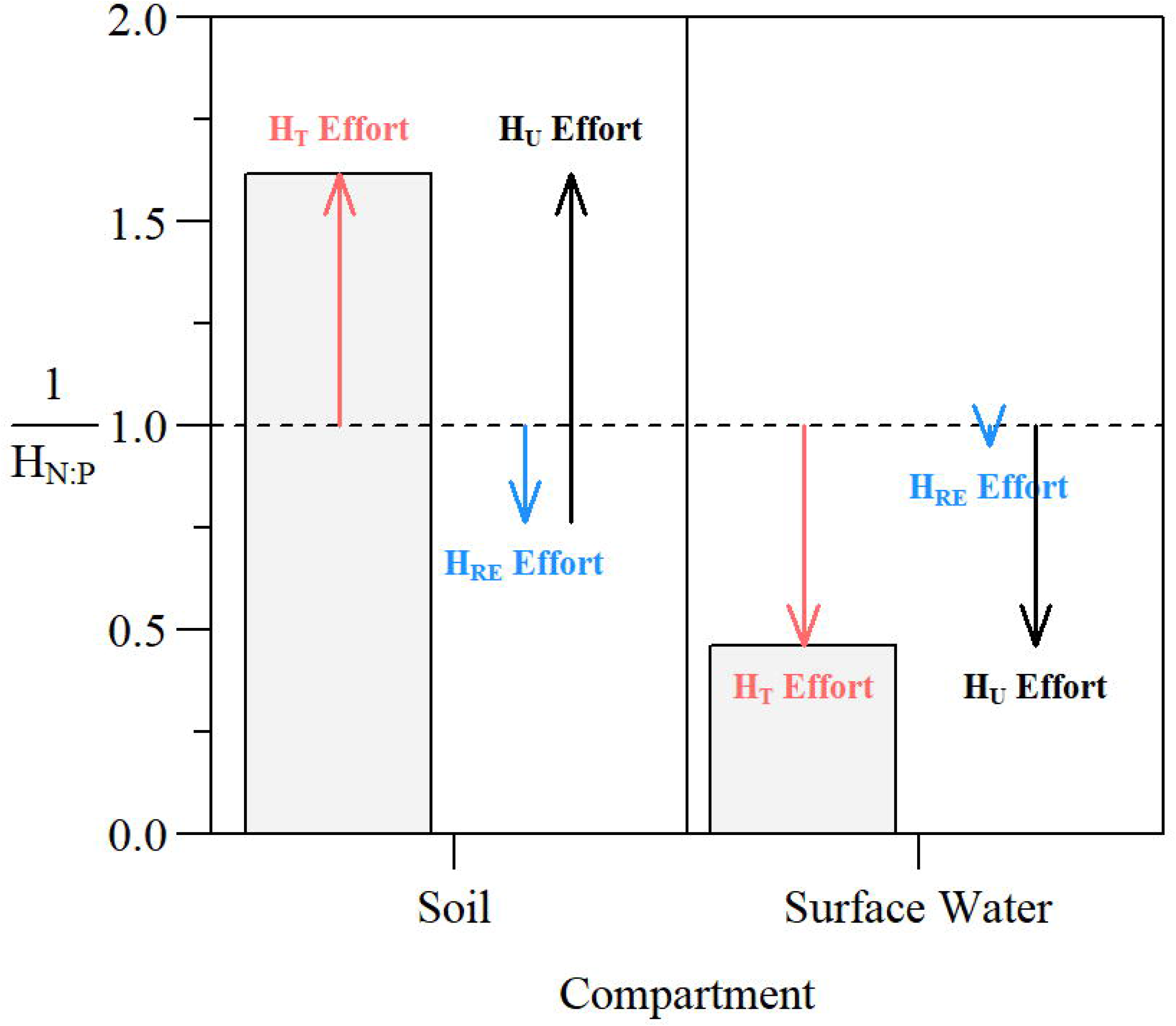
Median total homeostasis coefficient (1/*H*_*T*_) for emergent aquatic vegetation (EAV; FW1) with effort associated with maintaining total (*H*_*T*_; red arrows), resorptive (*H*_*RE*_; blue arrows) and uptake (*H*_*u*_; black arrows). A downward arrow indicates the effort into maintaining homeostasis, representing the 1-1/H expression in Eq. 5.

In addition to nutrient uptake, nutrient resorption itself can also contribute to overall stoichiometric homeostasis. In this study, we developed a concept based on first order dynamics similar to Menge et al. (2009) of how nutrient resorption can be incorporated into the stoichiometric homeostasis framework (Supplemental Material). By using equilibrium assumptions, we were able to distinguish contributions of uptake versus resorption to homeostasis (Fig. 6). In essence, we defined the effort put into homeostasis by the plant that determines changes in stoichiometry from differential uptake and from differential resorption. The definition of homeostasis is such that a value of zero indicates full homeostasis while a value of one suggests non-homeostasis. Effort towards homeostasis is therefore the complement of a homeostasis to one (Supplementary Information). The resulting calculations suggest that nutrient resorption contributes approximately 11% and 15% to stoichiometric homeostasis in EAV, when water and soil, respectively, are considered as the endmembers of nutrient supply (Fig. 6). In Fig.67 we indicated a positive effort towards maintaining homeostasis via an arrow down, with the length of the arrow indicating the magnitude of the effort. Due to the limitation of the data collected, we were unable to determine resorption homeostasis for SAV. Despite this limitation, given the resorption dynamics observed in EAV, it is possible that resorption homeostasis by SAV would be similar, contributing slightly to overall total homeostasis.

Despite some shortcomings of this study, the data allow us to evaluate homeostasic behavior in sub-tropical treatment wetland systems. As noted above, it is not clear to what degree the vegetation uses water versus soil as the main nutrient resource, generally it is assumed that EAV mine nutrients from the soil (via soil porewater) and SAV take up nutrients from the water column and modify the chemical environment to facilitate abiotic mineralization. It is possible that one nutrient is preferably obtained from the water column, and the other from the soil. Given the higher N:P in the soil relative to the water column, it would be opportune for plants to acquire N from the soil, while obtaining P from the water column. However, this inference may be complicated by the fact that vegetation is specifically mining soil P, depleting soil P storages and therefore increasing the N:P ratio in the soil. Given our data, it is not clear to what degree N or P are accessible for plant uptake. Along the FW, the proportion of inorganic (rapidly taken up) versus total nutrients varies in both the soil and water column (UF-WBL 2017). In the soil, the majority of nutrients could be in a mineral form (predominately Ca-P minerals) and therefore not as accessible to plants. Therefore, our calculation of homeostasis implicitly assumes that N and P were mineralized according to their total nutrient ratio. While not studied here, soil microbial interactions within the soil relative to ambient pool of nutrients could also influence site-specific nutrient dynamics along the FW (Wright and Reddy 2001; Corstanje et al. 2007). Inorganic nutrient addition experiments targeting soil and water column specifically, and perhaps at various levels and ratios, would further create a more nuanced picture with respect to homeostasis and preferential uptake pathways. Additionally, evaluation of SAV resorption and resorption specific to other plant parts including plant rhizomes (i.e. plant roots) would also fill knowledge gaps and allow better understanding of homeostasis due to resource partitioning within wetland macrophytes. Nevertheless, treatment wetlands are useful systems that can be used to study stoichiometric homeostasis as they present a wide nutrient gradient with large variability in nutrient loading from inflow to the outflow.

## Conclusions

This study suggests that vegetation within treatment wetlands maintains a relatively constant uptake efficiencies for nitrogen and phosphorus as indicated by the non-homeostatic relationship between foliar and active reservoirs. Thus, tissue nutrient concentrations and nutrient concentrations of the ambient environment scale with each other to some degree. In this study, we developed a framework to link nutrient resorption to the homeostasis concept and found that resorption efforts can alleviate nutrient imbalances in the resource pools, although the resource pool (water or soil) cannot be clearly identified. However, the strongly non-homeostatic behavior of EAV with respect to soil resources requires further attention and we speculate that a lowering of the N:P ratio in the resource pool will lead to storage of P in increasingly plant-inaccessible compounds.

## Acknowledgements

We would like to thank SFWMD and UF Wetland Biogeochemistry Laboratory staff members for providing the data used in this analysis. We would also like to thank Mark Brenner, Sue Newman, Odi Villapando, Tom James, Jim Elser and the anonymous peer reviewer(s) and editor(s) for their efforts and constructive review of this manuscript.

## Authors’ Contributions

PJ performed data analyses including necessary calculations and statistical analyses and wrote the manuscript. SG developed the expanded homeostasis framework, contributed to data interpretation and writing. RKB and TZO were involved with data collection and writing. ALW assisted with writing. JK was involved in data collection. All authors read and approved the final manuscript.

## References

Aerts R (1996) Nutrient Resorption from Senescing Leaves of Perennials: Are there General Patterns? Journal of Ecology 84:597–608. doi: 10.2307/2261481

Aerts R, Chapin FS (1999) The mineral nutrition of wild plants revisited: a re-evaluation of processes and patterns. Advances in ecological research 30:1–67

Boyer MLH, Wheeler BD (1989) Vegetation Patterns in Spring-Fed Calcareous Fens: Calcite Precipitation and Constraints on Fertility. Journal of Ecology 77:597–609. doi: 10.2307/2260772

Brix H (2003) Plants used in constructed wetlands and their functions. In: Dias V, Vymazel J (eds) 1st International Seminar on the use of Aquatic Macrophytes for Wastewater Treatment in Constructed Wetlands. Lisbon, Portugal

Bruland GL, Osborne TZ, Reddy KR, et al (2007) Recent Changes in Soil Total Phosphorus in the Everglades: Water Conservation Area 3. Environmental Monitoring and Assessment 129:379–395. doi: 10.1007/s10661-006-9371-x

Chapin III FS, Schulze E-D, Mooney HA (1990) The Ecology and Economics of Storage in Plants. Annual Review of Ecology and Systematics 21:423–447

Chen H, Ivanoff D, Pietro K (2015) Long-term phosphorus removal in the Everglades stormwater treatment areas of South Florida in the United States. Ecological Engineering 79:158–168. doi: 10.1016/j.ecoleng.2014.12.012

Chimney MJ, Pietro KC (2006) Decomposition of macrophyte litter in a subtropical constructed wetland in south Florida (USA). Ecological Engineering 27:301–321. doi: 10.1016/j.ecoleng.2006.05.016

Corstanje R, Reddy KR, Prenger JP, et al (2007) Soil microbial eco-physiological response to nutrient enrichment in a sub-tropical wetland. Ecological Indicators 7:277–289. doi: 10.1016/j.ecolind.2006.02.002

Craft CB, Vymazal J, Richardson CJ (1995) Response of everglades plant communities to nitrogen and phosphorus additions. Wetlands 15:258–271. doi: 10.1007/BF03160706

De Deyn GB, Cornelissen JHC, Bardgett RD (2008) Plant functional traits and soil carbon sequestration in contrasting biomes. Ecology Letters 11:516–531. doi: 10.1111/j.1461-0248.2008.01164.x

DeBusk TA, Kharbanda M, Jackson SD, et al (2011) Water, vegetation and sediment gradients in submerged aquatic vegetation mesocosms used for low-level phosphorus removal. Science of The Total Environment 409:5046–5056. doi: 10.1016/j.scitotenv.2011.08.038

Demers JD, Driscoll CT, Fahey TJ, Yavitt JB (2007) Mercury cycling in litter and soil in different forest types in the adirondack region, new york, usa. Ecological Applications 17:1341–1351. doi: 10.1890/06-1697.1

Dhote S, Dixit S (2009) Water quality improvement through macrophytes—a review. Environmental Monitoring and Assessment 152:149–153. doi: 10.1007/s10661-008-0303-9

Dierberg FE, DeBusk TA, Jackson SD, et al (2002) Submerged aquatic vegetation-based treatment wetlands for removing phosphorus from agricultural runoff: response to hydraulic and nutrient loading. Water research 36:1409–1422

Dinno A (2015) Dunn’s test of multiple comparisons using rank sums. CRAN R-Project Duarte CM (1992) Nutrient concentration of aquatic plants: Patterns across species. Limnol Oceanogr 37:882–889. doi: 10.4319/lo.1992.37.4.0882

Elser JJ, Fagan WF, Denno RF, et al (2000) Nutritional constraints in terrestrial and freshwater food webs. Nature 408:578

Elser JJ, Fagan WF, Kerkhoff AJ, et al (2010) Biological stoichiometry of plant production: metabolism, scaling and ecological response to global change. New Phytologist 186:593–608. doi: 10.1111/j.1469-8137.2010.03214.x

Feijoó C, Leggieri L, Ocón C, et al (2014) Stoichiometric homeostasis in the food web of a chronically nutrient-rich stream. Freshwater Science 33:820–831. doi: 10.1086/677056

Fernández-Aláez M, Fernández-Aláez C, Bécares E (1999) Nutrient content in macrophytes in Spanish shallow lakes. Hydrobiologia 408:317–326

Filipiak M (2016) Pollen Stoichiometry May Influence Detrital Terrestrial and Aquatic Food Webs. Front Ecol Evol 4:. doi: 10.3389/fevo.2016.00138

Filipiak M, Weiner J (2014) How to Make a Beetle Out of Wood: Multi-Elemental Stoichiometry of Wood Decay, Xylophagy and Fungivory. PLOS ONE 9:e115104. doi: 10.1371/journal.pone.0115104

Filipiak M, Weiner J (2017) Nutritional dynamics during the development of xylophagous beetles related to changes in the stoichiometry of 11 elements: Stoichiometry of xylophage development. Physiological Entomology 42:73–84. doi: 10.1111/phen.12168

Frost PC, Evans-White MA, Finkel ZV, et al (2005) Are you what you eat? Physiological constraints on organismal stoichiometry in an elementally imbalanced world. Oikos 109:18–28. doi: 10.1111/j.0030-1299.2005.14049.x

Gu Q, Zamin TJ, Grogan P (2017) Stoichiometric homeostasis: a test to predict tundra vascular plant species and community-level responses to climate change. Arctic Science 1–14. doi: 10.1139/as-2016-0032

Güsewell S (2004) N◻: P ratios in terrestrial plants: variation and functional significance. New Phytologist 164:243–266. doi: 10.1111/j.1469-8137.2004.01192.x

Güsewell S, Koerselman W (2002) Variation in nitrogen and phosphorus concentrations of wetland plants. Perspectives in Plant Ecology, Evolution and Systematics 5:37–61. doi: 10.1078/1433-8319-0000022

Güsewell S, Koerselman W, Verhoeven JTA (2003) Biomass n:p ratios as indicators of nutrient limitation for plant populations in wetlands. Ecological Applications 13:372–384. doi: 10.1890/1051-0761(2003)013[0372:BNRAIO]2.0.CO;2

Julian P, Gerber S, Bhomia RK, et al (2019) Evaluation of nutrient stoichiometric relationships among ecosystem compartments of a subtropical treatment wetland. Do we have “Redfield wetlands”? Ecol Process 8:20. doi: 10.1186/s13717-019-0172-x

Julian P, Gerber S, Wright AL, et al (2017) Carbon pool trends and dynamics within a subtropical peatland during long-term restoration. Ecol Process 6:43–57. doi: 10.1186/s13717-017-0110-8

Juston J, DeBusk TA (2006) Phosphorus mass load and outflow concentration relationships in stormwater treatment areas for Everglades restoration. Ecological Engineering 26:206–223. doi: 10.1016/j.ecoleng.2005.09.011

Kadlec RH, Wallace SD (2009) Treatment wetlands. CRC Press, Boca Raton, FL

Killingbeck KT (1996) Nutrients in Senesced Leaves: Keys to the Search for Potential Resorption and Resorption Proficiency. Ecology 77:1716–1727. doi: 10.2307/2265777

King J, Zweig C (2016) P Flux Vegetation Assessments-Submerged Aquatic Vegetation (SAV) Biomass Sampling and Analysis September 2016. South Florida Water Management District, West Palm Beach, FL

Koerselman W, Meuleman AFM (1996) The Vegetation N:P Ratio: a New Tool to Detect the Nature of Nutrient Limitation. The Journal of Applied Ecology 33:1441. doi: 10.2307/2404783

Komsta L (2013) Median-Based Linear Models. CRAN R-Project

Makino W, Cotner JB, Sterner RW, Elser JJ (2003) Are Bacteria More Like Plants or Animals? Growth Rate and Resource Dependence of Bacterial C: N: P Stoichiometry. Functional Ecology 17:121–130

McGroddy ME, Daufresne T, Hedin LO (2004) Scaling of C:N:P stoichiometry in forests worldwide: implications of terrestrial redfield-type ratios. Ecology 85:2390–2401. doi: 10.1890/03-0351

Menge DNL, Pacala SW, Hedin LO (2009) Emergence and Maintenance of Nutrient Limitation over Multiple Timescales in Terrestrial Ecosystems. The American Naturalist 173:164–175. doi: 10.1086/595749

Miao S (2004) Rhizome growth and nutrient resorption: mechanisms underlying the replacement of two clonal species in Florida Everglades. Aquatic Botany 78:55–66. doi: 10.1016/j.aquabot.2003.09.001

Miao SL, Zou CB (2012) Effects of inundation on growth and nutrient allocation of six major macrophytes in the Florida Everglades. Ecological Engineering 42:10–18. doi: 10.1016/j.ecoleng.2012.01.009

Newman S, Osborne TZ, Hagerthey SE, et al (2017) Drivers of landscape evolution: multiple regimes and their influence on carbon sequestration in a sub-tropical peatland. Ecol Monogr 87:578–599. doi: 10.1002/ecm.1269

Osborne TZ, Bruland GL, Newman S, et al (2011) Spatial distributions and eco-partitioning of soil biogeochemical properties in the Everglades National Park. Environmental Monitoring and Assessment 183:395–408. doi: 10.1007/s10661-011-1928-7

Peng H, Chen Y, Yan Z, Han W (2016) Stage-dependent stoichiometric homeostasis and responses of nutrient resorption in Amaranthus mangostanus to nitrogen and phosphorus addition. Scientific Reports 6:. doi: 10.1038/srep37219

Persson J, Fink P, Goto A, et al (2010) To be or not to be what you eat: regulation of stoichiometric homeostasis among autotrophs and heterotrophs. Oikos 119:741–751. doi: 10.1111/j.1600-0706.2009.18545.x

Rattray MR, Howard-Williams C, Brown JMA (1991) Sediment and water as sources of nitrogen and phosphorus for submerged rooted aquatic macrophytes. Aquatic Botany 40:225–237. doi: 10.1016/0304-3770(91)90060-I

Reddy KR, DeLaune RD (2008) Biogeochemistry of wetlands: science and applications. CRC Press, Boca Raton, FL

Reddy KR, Kadlec RH, Flaig E, Gale PM (1999) Phosphorus Retention in Streams and Wetlands: A Review. Critical Reviews in Environmental Science and Technology 29:83–146. doi: 10.1080/10643389991259182

Redfield AC (1958) The biological control of chemical factors in the environment. American Scientist 46:230A–221

Reed SC, Townsend AR, Davidson EA, Cleveland CC (2012) Stoichiometric patterns in foliar nutrient resorption across multiple scales. New Phytologist 196:173–180. doi: 10.1111/j.1469-8137.2012.04249.x

Rejmánková E (2005) Nutrient resorption in wetland macrophytes: comparison across several regions of different nutrient status. New Phytologist 167:471–482. doi: 10.1111/j.1469-8137.2005.01449.x

Rejmánková E, Macek P, Epps K (2008) Wetland ecosystem changes after three years of phosphorus addition. Wetlands 28:914–927. doi: 10.1672/07-150.1

Rejmánková E, Snyder JM (2008) Emergent macrophytes in phosphorus limited marshes: do phosphorus usage strategies change after nutrient addition? Plant Soil 313:141–153. doi: 10.1007/s11104-008-9687-0

Richardson CJ, Marshall PE (1986) Processes Controlling Movement, Storage, and Export of Phosphorus in a Fen Peatland. Ecological Monographs 56:280–302. doi: 10.2307/1942548

Rivas-Ubach A, Sardans J, Pérez-Trujillo M, et al (2012) Strong relationship between elemental stoichiometry and metabolome in plants. PNAS 109:4181–4186. doi: 10.1073/pnas.1116092109

Sardans J, Rivas-Ubach A, Peñuelas J (2012) The C:N:P stoichiometry of organisms and ecosystems in a changing world: A review and perspectives. Perspectives in Plant Ecology, Evolution and Systematics 14:33–47. doi: 10.1016/j.ppees.2011.08.002

Shaver GR, Melillo JM (1984) Nutrient Budgets of Marsh Plants: Efficiency Concepts and Relation to Availability. Ecology 65:1491–1510. doi: 10.2307/1939129

Sistla SA, Appling AP, Lewandowska AM, et al (2015) Stoichiometric flexibility in response to fertilization along gradients of environmental and organismal nutrient richness. Oikos 124:949–959. doi: 10.1111/oik.02385

Sistla SA, Schimel JP (2012) Stoichiometric flexibility as a regulator of carbon and nutrient cycling in terrestrial ecosystems under change. The New Phytologist 196:68–78

Small GE, Helton AM, Kazanci C (2009) Can consumer stoichiometric regulation control nutrient spiraling in streams? Journal of the North American Benthological Society 28:747–765. doi: 10.1899/08-099.1

Smalley GW, Coats DW, Stoecker DK (2003) Feeding in the mixotrophic dinoflagellate Ceratium furca is influenced by intracellular nutrient concentrations. Marine Ecology Progress Series 262:137–151. doi: 10.3354/meps262137

South Florida Water Management District (2013) Restoration Strategies Regional Water Quality Plan: Science Plan for the Everglades Stormwater Treatment Areas. South Florida Water Management District, West Palm Beach, FL

Sterner, Clasen, Lampert, Weisse (1998) Carbon:phosphorus stoichiometry and food chain production. Ecology Letters 1:146–150. doi: 10.1046/j.1461-0248.1998.00030.x

Sterner RW, Elser JJ (2002) Ecological Stoichiometry: The Biology of Elements from Molecules to the Biosphere. Princeton University Press

UF-WBL (2017) Evaluation of Soil Biogeochemical Properties Influencing Phosphorus Flux in the Everglades Stormwater Treatment areas: 2016-2017 Annual Report. University of Florida, Gainesville, FL

Van Heerwaarden LM, Toet S, Aerts R (2003) Current measures of nutrient resorption efficiency lead to a substantial underestimation of real resorption efficiency: facts and solutions. Oikos 101:664–669. doi: 10.1034/j.1600-0706.2003.12351.x

Villapando O, King J (2018) Appendix 5C-3: Evaluation of Phosphorus Sources, Forms, Flux, and Transformation Processes in the Stormwater Treatment Areas. In: 2018 South Florida Environmental Report. South Florida Water Management District, West Palm Beach, FL

Wright AL, Reddy KR (2001) Phosphorus loading effects on extracellular enzyme activity in Everglades wetland soils. Soil Science Society of America Journal 65:588–595

Yu Q, Elser JJ, He N, et al (2011) Stoichiometric homeostasis of vascular plants in the Inner Mongolia grassland. Oecologia 166:1–10. doi: 10.1007/s00442-010-1902-z

Zar JH (2010) Biostatistical Analysis, 15th edn. Prentice Hall

