## Supplemental Material for "Stoichiometric homeostasis of wetland vegetation along a nutrient gradient in a subtropical wetland. Understanding stoichiometric mechanisms of nutrient retention in wetland macrophytes"

Stoichiometric homeostasis is estimated by the equation below (Eq. S1) consistent with Eq. 2 of the main text.

$$\frac{1}{H_{T}}=\frac{\log\left( y \right)-\log\left( c \right)}{\log\left( x \right)}$$

Eq. S1

Where $y$ represents the consumer stoichiometry, $x$ represents the resource stoichiometry $c$ is a constant determined from the log-log regression of consumer and resource stoichiometry, and *H_T_* is the total homeostasis. This metric provides a homeostasis coefficient that relates consumer stoichiometry to the resource stoichiometry. Building from this concept, we develop relationships to quantify the consumer’s effort to maintain homeostasis by two pathways, differential uptake and differential resorption, i.e. a dynamic representation of homeostasis.

Using very simple first order dynamics for consumer (similar to Menge et al. 2009), the nutrient dynamics of an organism can be expressed with

$$\frac{dY_{N}}{dt}=a_{N}\times X_{N}-k\times Y_{N}\times(1-RE_{TN})$$

Eq. S2

$$\frac{dY_{P}}{dt}=a_{P}\times X_{P}-k\times Y_{P}\times(1-{RE_{TP}})$$

Eq. S3

Where *Y_N_* and *X_N_* is the nitrogen concentration of the consumer and resource, respectively, *Y_P_* and *Y_P_* the phosphorus concentration of the two groups, *a_N_* and *a_P_* are the uptake coefficients, *k* the tissue turnover rate of the consumer, and *RE­_TN_* and *RE_TP_* are the resorption coefficients.

At quasi steady state (i.e. no growth, $\frac{dY_{N}}{dt}=\frac{dY_{P}}{dt}=0$), the stoichiometric ratio becomes:

| $\frac{Y_{N}}{Y_{P}}=\frac{a_{N}}{a_{P}}\times\frac{X_{N}}{X_{P}}\times\frac{\left( 1-RE_{Tp} \right)}{\left( 1-{RE_{TN}} \right)}$=$c\left( \frac{X_{N}}{X_{P}} \right)^{1/H_{T}}$ | Eq. S4 |
| --- | --- |

The expression on the right hand side is derived from taking the exponent on both sides of Eq. S1 and then solve for *y* (which is the stoichiometric ratio of the consumer). Dividing both sides of Eq S4 by *X_N_/X_P_*, we find

$$\frac{a_{N}}{a_{P}}\times\frac{\left( 1-RE_{TP} \right)}{\left( 1-RE_{TN} \right)}= c\left( \frac{X_{N}}{X_{P}} \right)^{\frac{1}{H_{T}}-1}$$

Eq. S5

which is and expression of how uptake (a) and resorption are linked to the homeostasis coefficient. The two factors can be separated to obtain their own homeostasis coefficient. The contribution of uptake to homeostasis is::

$$\frac{a_{N}}{a_{P}}= c_{U}\left( \frac{X_{N}}{X_{P}} \right)^{\frac{1}{H_{U}}-1}$$

Eq. S6

Where the term 1/H_U_ is the uptake homeostasis coefficient. Likewise, we have:

$$\frac{\left( 1-r_{P} \right)}{\left( 1-r_{N} \right)}= c_{R}\left( \frac{X_{N}}{X_{P}} \right)^{\frac{1}{H_{R}}-1}$$

Eq. S7

For the resorption’s contribution (H_R_).

Substituting Eq. S6 and S7 into S4

$$\frac{Y_{N}}{Y_{P}}=c_{U}\left( \frac{X_{N}}{X_{P}} \right)^{\frac{1}{H_{U}}-1}c_{R}\left( \frac{X_{N}}{X_{P}} \right)^{\frac{1}{H_{R}}-1}\frac{X_{N}}{X_{P}}={c\left( \frac{X_{N}}{X_{P}} \right)}^{1/H_{T}}$$

Eq. S8

Taking the log yields

$$\log\left( c \right)+\frac{1}{H_{T}}*\log\left( x \right)=\log\left( c_{U} \right)+\log\left( c_{R} \right)+\left( \frac{1}{H_{U}}-1+\frac{1}{H_{R}}-1+1 \right)*log(x)$$

Eq. S9

Where *x=X_N_/X_P_* and *y=Y_N_/Y_P_* are the ratios of the nutrient in the resource and consumer respectively)

Rearranging:

$$\log\left( c \right)+\left( \frac{1}{H_{T}}-1 \right)\log\left( x \right)=\left( \frac{1}{H_{U}}-1 \right)\log\left( x \right)+\left( \frac{1}{H_{R}}-1 \right)\log\left( x \right)+\log\left( c_{U} \right)+log(c_{R})$$

Eq. S10

*H_T_* and *H_R_* are quantities obtained from slopes regressing organism ratio and resorption ratios, respectively, against resource ratio, which allow solving for the unknown *H_U_*:

$$\left( \frac{1}{H_{T}}-1 \right)=\left( \frac{1}{H_{U}}-1 \right)+\left( \frac{1}{H_{R}}-1 \right)$$

Eq. S11

(Note, that the regression constants drop if *c*:*= c_U_*c_R_*). By multiplying the equation above with (-1):

$$\left( 1-\frac{1}{H_{T}} \right)=\left( 1-\frac{1}{H_{U}} \right)+\left( 1-\frac{1}{H_{R}} \right)$$

Eq. S12

Where the expression $\frac{1}{H_{i}}-1$ (i=T,U,R) can be viewed as *effort* towards maintaining homeostasis by the entire plant (T), partitioned into its component uptake (U) and resorption (R).

**References**

Menge DNL, Pacala SW, Hedin LO (2009) Emergence and Maintenance of Nutrient Limitation over Multiple Timescales in Terrestrial Ecosystems. Am Nat 173:164–175. doi: 10.1086/595749
