## Supplementary material for "Stoichiometric homeostasis of wetland vegetation along a nutrient gradient in a subtropical wetland. Understanding stoichiometric mechanisms of nutrient retention in wetland macrophytes": Table S1

Table S1. Summary of parameters, matrices, analytical methods and minimum detection limit (MDL) used for this study. Additional parameters were collected but not used in this study. All analytical methods are consistent with Florida Department of Environmental Protection or U.S. Environmental Protection Agency Standard Operating Procedures and methods.

| **Matrix** | **Parameter** | **Abbreviation** | **Analytical Method** | **Minimum Detection Limit** | **Method** |
| --- | --- | --- | --- | --- | --- |
| Surface Water | Total Phosphorus | TP | SM4500PF | 2 µg P L^-1^ | Clesceri et al. (1998) |
|  | Total Nitrogen | TN | SM4500NC | 0.02 mg N L^-1^ | Clesceri et al. (1998) |
|  | Dissolved Organic Carbon | DOC | SM5310B | 0.8 mg C L^-1^ | Clesceri et al. (1998) |
| Soil and Vegetation | Loss-on-ignition^1,2^ | LOI | Calculation^2^ | 1.0 % | **---** |
|  | Total Phosphorus | TP | SM4500PF | 16 mg P kg^-1^ | Clesceri et al. (1998) |
|  | Total Nitrogen | TN | SFWMD 3200 | 2 g N kg^-1^ | SFWMD (2015) |
|  | Total Carbon | TC | SFWMD 3200 | 2 g C kg^-1^ | SFWMD (2015) |
|  | Total Calcium^1^ | TCa | EPA 6010C | 109 mg Ca kg^-1^ | US EPA (2007) |

^1^ Loss-on-ignition and total calcium was assessed for soil components only.

^2^ Loss-on-ignition was calculated from the difference between 100% and percent ash determined by the analytical method identified as SFWMD 1610 (SFWMD 2015)**.**
